# LinearCapR: Linear-time computation of per-nucleotide structural-context probabilities of RNA without base-pair span limits

**DOI:** 10.64898/2025.12.26.696559

**Authors:** Takumi Otagaki, Hiroaki Hosokawa, Tsukasa Fukunaga, Junichi Iwakiri, Goro Terai, Kiyoshi Asai

## Abstract

**Motivation:** RNA molecules adopt dynamic ensembles of secondary structures, where the local structural context of each nucleotide–such as whether it resides in a stem or a specific type of loop–strongly shapes molecular interactions and regulatory function. Structural-context probabilities therefore provide a more functionally informative view of RNA folding than the minimum free energy structures or base-pairing probabilities. However, existing tools either require *O* (*N*^3^) time or employ span-restricted approximations that omit long-range base-pairs, limiting their applicability to large and biologically important RNAs.

**Results:** We introduce LinearCapR, enabling linear-time, span-unrestricted computation of structural-context marginalized probabilities, using beam-pruned Stochastic Context Free Grammar-based computation. LinearCapR retains global ensemble features lost by span-limited methods and yields superior predictive power on bpRNA-1m(90) dataset, especially for multiloops and exterior regions, as well as long-distance stems. LinearCapR supports analysis of long RNAs, demonstrated on the full genome of SARS-CoV-2.

**Conclusions:** LinearCapR provides the first base-pair-span-unrestricted, linear-time framework for RNA structural-context analysis, retaining key thermodynamic ensemble features essential for functional interpretation. It enables large-scale studies of viral genomes, long non-coding RNAs, and downstream analyses such as RNA-binding protein site prediction.

**Availability:** The source code of LinearCapR is available at https://github.com/hoget157/LinearCapR.

## 1 Introduction

RNA molecules form highly dynamic ensembles of secondary structures rather than a single static conformation. Their conformational states fluctuate according to thermodynamic stability and environmental conditions, making both the determination and prediction of RNA structures a challenging problem. While high-resolution experimental approaches such as NMR spectroscopy, X-ray crystallography, and cryo-electron microscopy can reveal detailed architectures, they are labor- and cost-intensive and do not readily capture alternative conformations at scale.

Computational methods based on nearest-neighbor thermodynamic models provide an efficient alternative. Under pseudoknot-free assumption, dynamic programming enables the computation of the minimum free energy (MFE) structures [Zuker, 1989] or partition functions and base-pairing probabilities (BPPs) [McCaskill, 1990] in *O* (*N*^3^) time and *O* (*N*^2^) space.The ensemble-level quantities (partition functions and BPPs) offer essential insight into alternative conformations, regulatory switching, and structural accessibility that underlie RNA functions.

However, base-pairing information alone is insufficient for many biological questions. The structural context of each nucleotide — such as whether it is located in a stem, hairpin loop, bulge, internal loop, multiloop, or exterior region —strongly influences RNA-binding proteins, catalytic activity of ribozymes, and post-transcriptional regulation. Posterior probabilities over these structural contexts therefore provide a richer and more functionally meaningful representation of RNA structural ensemble than BPPs alone.

CapR introduced a stochastic context free grammar (SCFG)-based framework to compute such structural-context probabilities for each nucleotide [Fukunaga et al., 2014], but its original formulation requires *O* (*N*^3^) time, limiting its applicability to long RNAs. To improve scalability, CapR and related tools such as RNAplfold and Rfold impose a base-pair span limit *W*, reducing computational cost to *O* (*NW* ^2^) [Bernhart et al., 2006, Kiryu et al., 2008]. Yet span limit necessarily discards long-range base-pairing interactions, which are widespread in natural RNAs – for example, compact viral genomes and end-to-end contacts in mRNAs [Yoffe et al., 2011, Lai et al., 2018]. Thus, efficient and span-unrestricted context profiling has remained elusive.

Beam-search-based linearization provides an effective means of approximating SCFG-based computation in linear time by pruning low-probability states during left-to-right parsing. While related pruning strategies have been explored in other grammar-driven domains [Chiang, 2007], the breakthroughs most relevant to RNA biology are LinearFold [Huang et al., 2019] and LinearPartition [Zhang et al., 2020], whose beam-search formulations established that global RNA folding statistics can be approximated with remarkable speed and accuracy. Their success provided a new computational paradigm for RNA structure prediction. Inspired by these developments, we bring beam-pruned computation to a different but complementary objective: computing nucleotide-level structural-context posterior probabilities, which has not previously been addressed by any span-unrestricted or beam-based method.

Here, we introduce LinearCapR, the first linear-time algorithm for computing CapR-style structural-context probabilities without imposing a base-pair span limit. LinearCapR reformulates CapRâĂŹs SCFG-based computation within a beam-pruning framework inspired by LinearFold and LinearPartition, enabling efficient inside-outside recursion while preserving long-range base-pairing interactions.

Beam pruning bounds the number of candidate states but does not by itself linearize structural-context computation. LinearCapR therefore introduces constant-time range updates for structural-context profiling, removing residual length dependence and enabling end-to-end linear-time computation.

This advance enables accurate and scalable transcriptome- and genome-wide structural-context analysis, supporting downstream studies that benefit from interpretable, ensemble-aware RNA structure representations.

## 2 Materials and Methods

### 2.1 Algorithm

#### 2.1.1 Overview of LinearCapR

LinearCapR approximates CapR’s probabilistic ensemble using a beam-pruned inside-outside dynamic program that scans the sequence from 5^′^ to 3^′^. It maintains at most *b* highest-weighted partial structures (“states”) per DP node, pruning low-weight states for reducing search complexity.

CapR linearizes its dynamic programming with time complexity *O* (*NW* ^2^) by restricting maximum span of base-pairs to a fixed value *W* for an RNA sequence of length *N*. Since LinearCapR imposes no explicit maximum span on base pairs, long-range interactions remain accessible. For a fixed beam width *b* and unpaired-run cap *C*, LinearCapR achieves: **Time:** *O* (*Nb*^2^ + *NbC*^2^), **Space:** *O* (*Nb*).

#### 2.1.2 Problem Formulation

Consider an RNA sequence of length *N* indexed by *i* = 1, …, *N*. We restrict secondary structures to pseudoknot-free conformations. Each nucleotide *i* belongs to exactly one of the following six structural contexts:

- Stem (S): forms a base pair
- Hairpin loop (H): enclosed by a single base pair
- Bulge (B): in a two-base-pair loop with one-sided unpaired run
- Internal loop (I): in a two-base-pair loop with two-sided unpaired run
- Multiloop (M): enclosed by ≥ 3 closing base pairs
- Exterior loop (E): not enclosed by any base pair

Figure 1 illustrates how each nucleotide in the example secondary structure falls into one of the six context categories. The **structural-context profile** is the set of posterior probabilities: *p* (*i, δ*) ≡ *Pr* [*x*_*i*_ is in context *δ*], where *δ* ∈ {S, H, B, I, M, E}.

**Figure 1.**
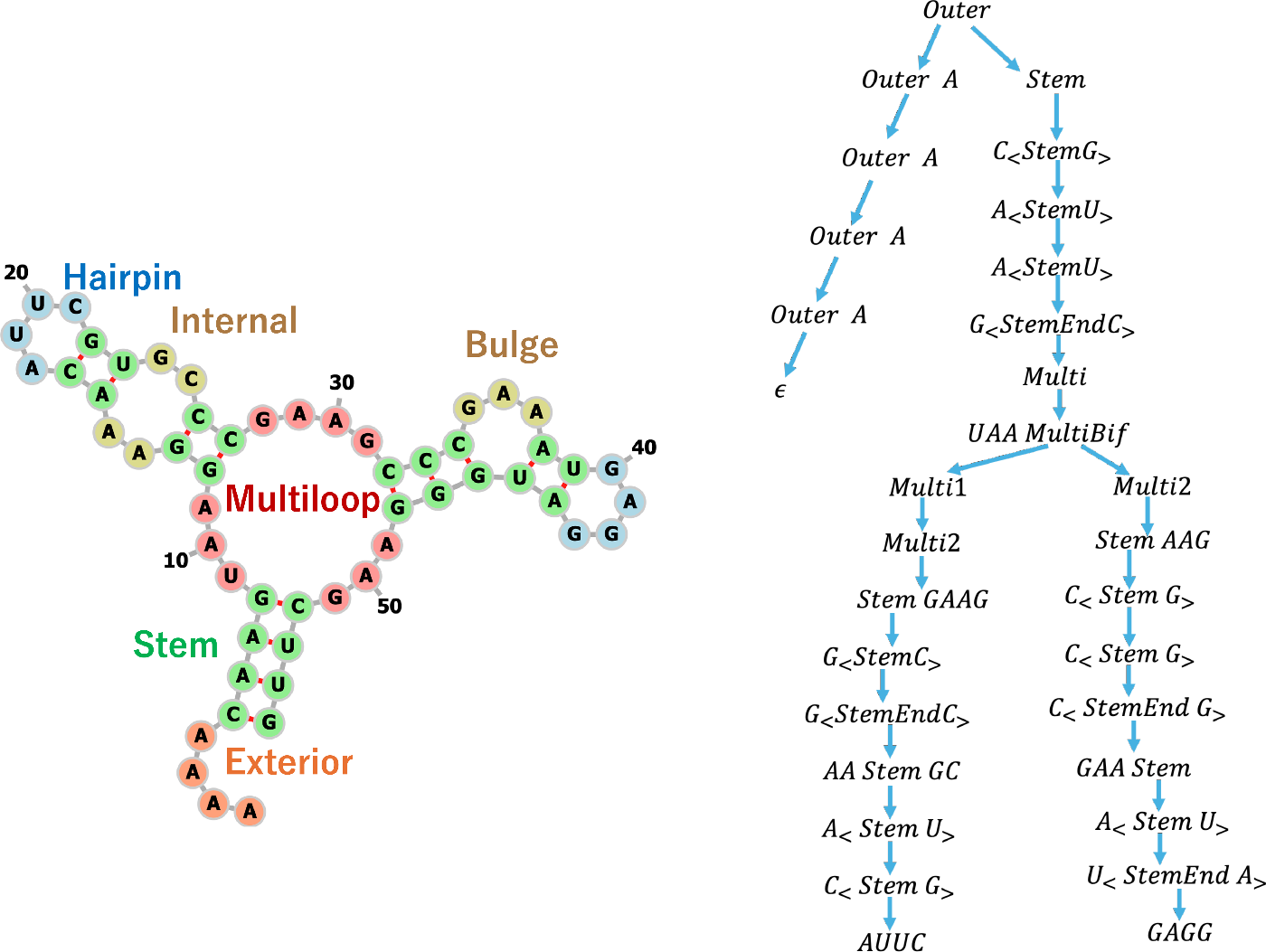
Left: Representative RNA secondary structure annotated with the six structural contexts: stem (S), hairpin (H), bulge (B), internal (I), multiloop (M), and exterior (E). The figure was drawn by Forna [Kerpedjiev et al., 2015]. Right: Example derivation under the stochastic context-free grammar introduced in the Materials and Methods 2.1.3 (The Grammar and the Algorithm), illustrating how the start symbol Outer is expanded via the Stem, StemEnd, and multiloop production rules to generate the pseudoknot-free structure shown on the left.

#### 2.1.3 The Grammar and the Algorithm

We adopt the stochastic context-free grammar (SCFG) used in CapR [Fukunaga et al., 2014] to generate pseudoknot-free RNA secondary structures. The grammar consists of the following seven non-terminal symbols summarized in Table 1.

**Table 1:** Non-terminal symbols used in the SCFG.

| Name | Description |
| --- | --- |
| **Outer** | Exterior unpaired region |
| **Stem** | Helix extension |
| **StemEnd** | Terminates a helix into a hairpin, bulge/internal loop, or multiloop region |
| **Multi** | Completed multiloop |
| **MultiBif** | Bifurcation into two or more branches |
| **Multi1** | One-or-more list of branches |
| **Multi2** | One branch followed by a 3'-side run of unpaired bases |

The production rules are:

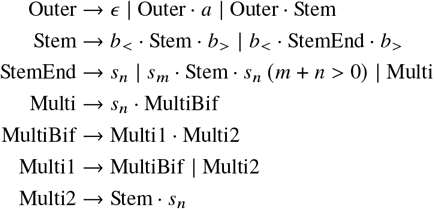

where *ϵ* is the empty string, *a* is a single unpaired nucleotide, *b*_*<*_, *b*_*>*_ denote the left/right sides of a base pair, and *s*_*n*_ represents a run of *n* consecutive unpaired nucleotides. Here, the dot “·” denotes concatenation of symbols. This decomposition mirrors the factorization required by the Inside recursion. The pseudo-code of the algorithm is shown in Algorithm 1, where *t* (*A → B*) denotes the Boltzmann weight of production *A* → *B,α*_*s*_ (*i, j*) (*β*_*s*_ (*i, j*)) stores the Inside (Outside) score of state *s* spanning indices *i* through *j* . The corresponding Outside pass is provided in Supplementary S1 Algorithm.

To enable linear-time pruning we have removed the recursive productions on `Multi` and `Multi2` from the SCFG of Rfold, and bound the number of consecutive unpaired bases by a constant *C* . We impose *n* ≤ *C* and *m* + *n* ≤ *C* to limit bulge/internal loop sizes and the number of single-strand residues between multibranch helices. Following CapR we set *C* = 30 in all the experiments. Relative to the Rfold grammar, removing the recursion from `Multi` and `Multi2` forces multiloop unpaired segments to be generated directly and shrinks the multiloop recurrences from *O* (*N*) candidate transitions to *O* (*C*).

Any pseudoknot-free structure whose unpaired runs do not exceed *C* can be generated by this grammar through repeated expansion starting from Outer. Therefore, the grammar provides a complete search space for commonly observed conformations while bounding the fan-out that the beam search needs to explore. The *C* truncation makes the grammar an approximation for the rare loops that exceed 30 contiguous unpaired bases (e.g., when a hairpin, internal, bulge or multiloop consecutive unpaired segment spans tens of nucleotides on one side), but in the bpRNA multiloop regions 99.57% of unpaired runs are 30 nt or shorter (Supplementary Fig. S1), so the approximation leaves only 0.43% of multiloop unpaired stretches outside the search space.

#### 2.1.4 Energy Models

We assume that the probability of an RNA adopting a secondary structure *σ* follows a Boltzmann distribution proportional to exp (−*G* (*σ*) / (*RT*)), where *G* (*σ*) is the free-energy of *σ, R* is the gas constant (1.98717 cal/K), and *T* is set to 310.15 K (37 °C). Loop energies are evaluated with the Turner2004 or Turner1999 nearest-neighbour parameters [Mathews et al., 2004] in this paper. Non-canonical base pairs present in the datasets are treated as unpaired. LinearCapR supports both Turner2004 (default) and Turner1999 parameters via a command-line option. Unless otherwise noted, we report results with Turner2004.

#### 2.1.5 Calculation of structure profile

During the Inside pass, we process sequence positions from 1 to *N* and retain only the top-*b* states for each non-terminal at the current frontier before distributing their scores to successor states.

The pruning threshold is computed by a linear-time selection routine, similar to QuickSelect [Randell and Russell, 1963], so that the expected cost of pruning is linear in the beam size. As a result, the overall inside computation runs in *O* (*Nb*^2^ + *NbC*^2^) time and *O* (*Nb*) space when the *b* and the *C* are treated as constants.

To avoid numerical underflow, the implementation accumulates values in log-space and applies a polynomial approximation of `log_sum_exp` following prior work in CONTRAfold [Do et al., 2006] and LinearPartition[Zhang et al., 2020].

The Outside recursion is performed in reverse order over the surviving Inside states: for each state *s*, we compute *β*_*s*_ (*i, j*) only for pairs (*i, j*) that survived the Inside recursion, which keeps the Outside complexity asymptotically the same as the Inside pass. Algorithm 1 shows the Inside pass, while the complete Outside recursion and full probability aggregation (hairpin, bulge/internal, stem, exterior) are provided in Supplementary Algorithm S1.

Beam pruning alone is not sufficient to fully linearize CapR-style structural-context computation, because structural-context probabilities must be accumulated over contiguous nucleotide ranges for each surviving state. To remove this remaining source of length-dependent cost, LinearCapR introduces constant-time range updates for profile aggregation. Using difference-array buffers, range contributions are recorded without updating every nucleotide position, and the final profiles are recovered by a single prefix-sum pass per context. This avoids explicit per-position updates within each range and keeps the cost of profile aggregation independent of range length, allowing the overall computation to scale effectively linearly.

Algorithm 2 illustrates this strategy for the multiloop case. We use *B*_MultiUnpaired_ to denote the Boltzmann-weighted contributions of multiloop unpaired segments. The `Multi` non-terminal tracks the unpaired segments between the loop’s branches, so *p* ([ *i, p* − 1 ], *M*) is added only for the unpaired nucleotides, even though the loop is closed by multiple stems. The resulting structure profile satisfies ∑_*δ*_ *p* (*i, δ*) = 1 for every position *i* and provides the desired marginal probabilities for each nucleotide and structural context.

##### Algorithm 1

Algorithm for the inside algorithm

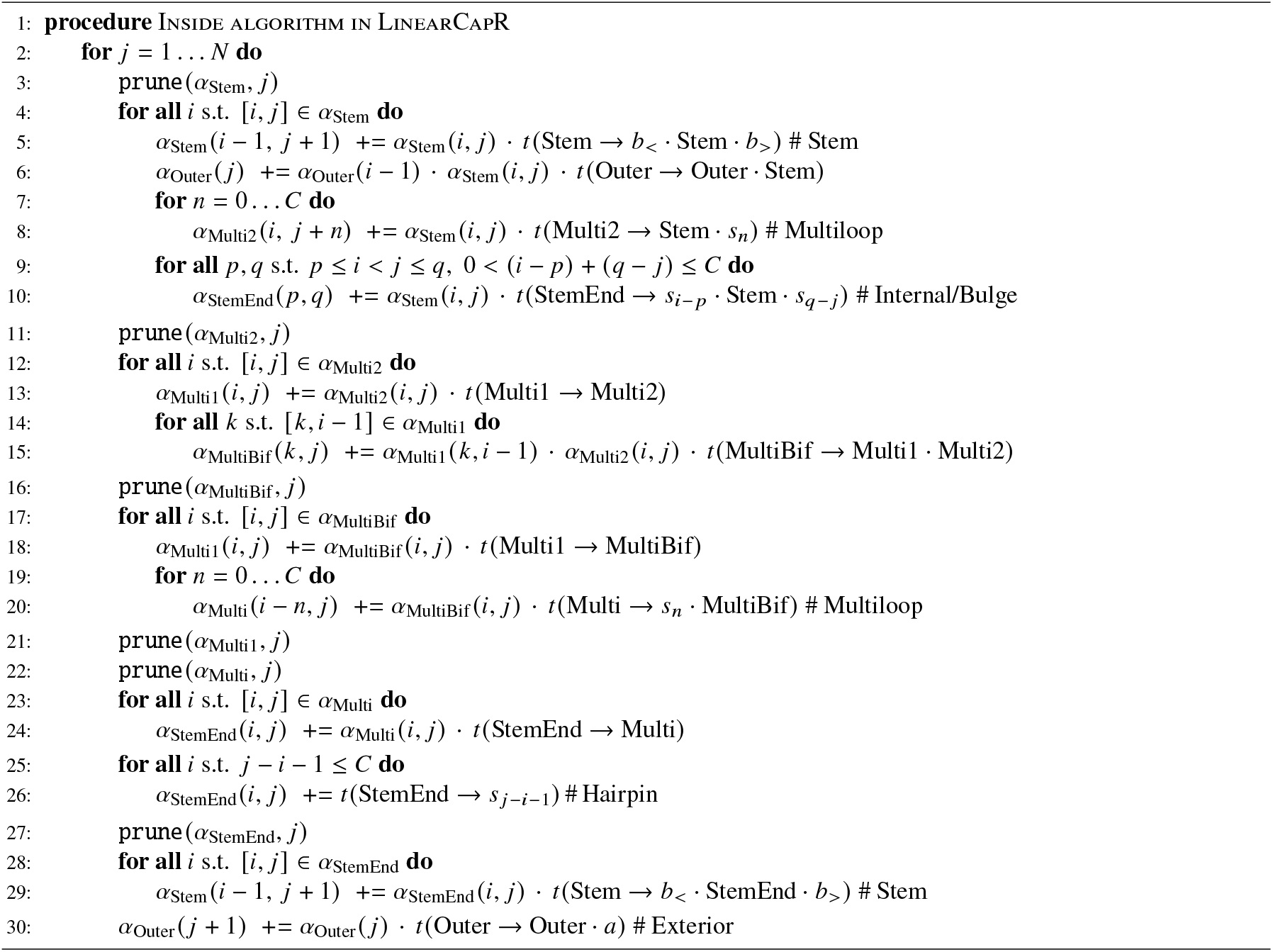

##### Algorithm 2

Multiloop contributions to the structure profile

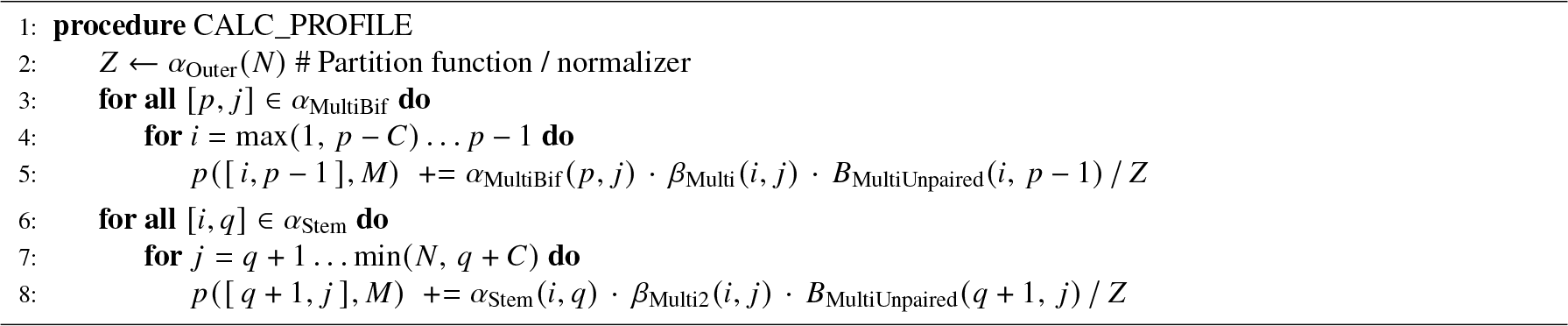

The full Outside recursion and the remaining probability updates (hairpin, bulge/internal, stem, exterior) are provided in Supplementary Text S1.

### 2.2 Dataset

1. **bpRNA-1m(90)**: We extracted pseudoknot-free sequences from the bpRNA-1m(90) database[Danaee et al., 2018] by selecting entries that belong to Rfam families [Griffiths-Jones et al., 2003] and whose annotated structures have page number ≤ 1 [Clote et al., 2012], a criterion that excludes pseudoknots. The resulting subset contains 24,901 sequences with lengths ranging from 11 to 4,065 nt, and each with an RNA sequence and an associated reference secondary structure for accuracy evaluation.
2. **RNAcentral**: To stress-test performance on long RNAs, we selected 20 sequences from RNAcentral [RNAcentral Consortium, 2021] covering a broad length range (4,000–985,945 nt). We divided this range into 20 logarithmically spaced bins and randomly sampled one sequence from each bin (see Supplementary S2 for exact bin edges).
3. **SARS-CoV-2 genome**: We further analyzed the 29,903 nt positive-sense RNA genome of SARS-CoV-2 (accession NC_045512)[Rangan et al., 2020] to examine how beam width affects performance on a single, moderately long viral genome.

### 2.3 Computational Experiments

We designed three complementary sets of experiments to assess (i) runtime and memory usage using bpRNA-1m(90) and RNAcentral datasets, (ii) the accuracy of the structural-context profiles using bpRNA-1m(90), and (iii) the effect of beam width on the partition-function approximation using SARS-CoV-2 genome.

For the accuracy analysis, we obtained ground-truth labels for the six structural contexts by parsing each bpRNA-1m(90) structure with the same SCFG-inspired decomposition used by CapR[Danaee et al., 2018]. Because we restrict to pseudoknot-free entries, every nucleotide maps unambiguously to exactly one context class, which provides a well-defined basis for the ROC/AUC evaluations.

Benchmark dataset sizes and length ranges are summarized in Supplementary Table S1.

#### 2.3.1 Execution Environment

All experiments (bpRNA, RNAcentral, SARS-CoV-2) were run on the same machine: dual-socket Intel Xeon Platinum 8360Y CPUs (72 hardware threads, 2.40 GHz) with 503 GiB RAM under Linux 4.18. The LinearCapR source code was compiled with a GNU C++17 compiler using `-O3 -std=c++17 -Wall`. No GPU acceleration or multi-node distribution was used.

## 3 Results

### 3.1 Run time and memory usage

To quantify scalability, we measured wall-clock runtime and peak memory usage as a function of sequence length on the bpRNA-1m(90) and on a panel of long RNAs from RNAcentral. Because the time complexity of CapR and LinearCapR depends quadratically on the span limit *W* or beam width *b* (CapR: (*O* (*NW* ^2^); LinearCapR: *O* (*Nb*^2^ *N* + *bC*^2^))), we varied these parameters over the same range, *b, W* ∈ { 50, 100, 200 }, and recorded runtime and peak memory for each sequence.

For both methods, runtime and memory increased linearly with length once *W* or *b* was fixed (Fig.2, left), consistent with the expected *O* (*N*) scaling. Runtime differences were modest: CapR exhibits lower runtime (e.g. 7.2s vs. 12.5s at 4,000nt when *b* = *W* = 100) due to its simpler state representation, whereas LinearCapR trades modest additional overhead for span-unrestricted ensemble fidelity. Memory usage showed a clear distinction: LinearCapR retains more active states, resulting in steeper memory growth (3.9GB vs. 1.6GB at *b* = *W* = 100).

**Figure 2.**
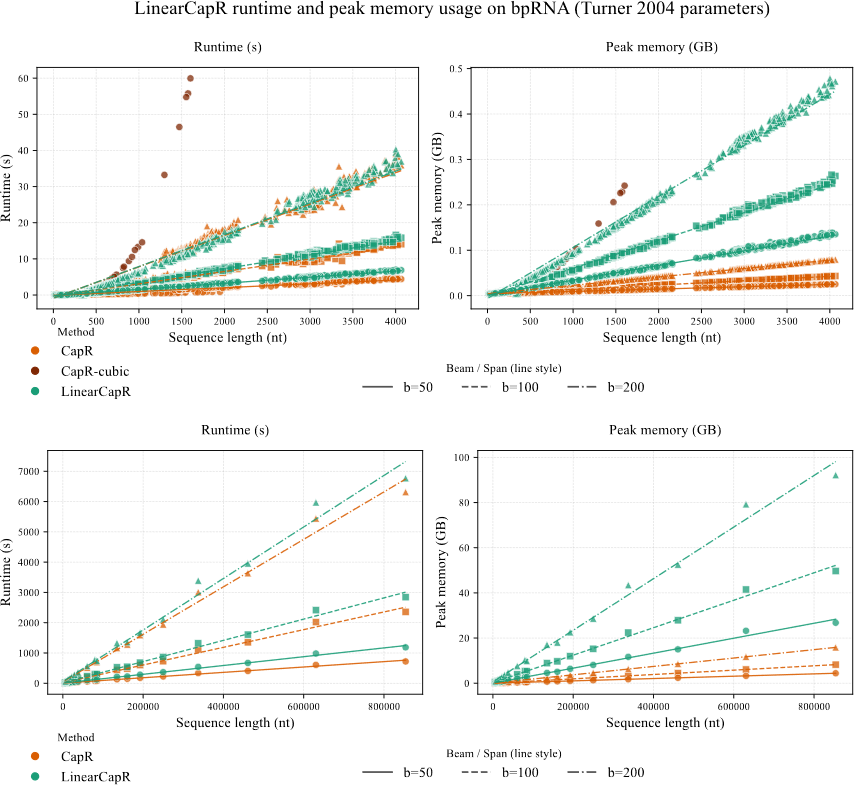
Runtime (top) and peak memory (bottom) versus sequence length for LinearCapR (beam widths *b* ∈ {50, 100, 200}), CapR (span limit *W* ∈ {50, 100, 200}) and CapR-cubic (original cubic-time CapR implementation without span limits). Solid lines denote bpRNA-1m(90) (left panel) and RNAcentral (right panel); each line reflects the per-parameter linear regression fits (one regression per beam width/span limit). CapR-cubic denotes the original *O* (*N*^3^) implementation without span constraints, which is impractical for long sequences but is included here as an upper-bound benchmark.

We next evaluated performance on long RNAcentral sequences ranging from 4kb to 853kb (Fig.2, right). Linear scaling was preserved across three order of magnitude in sequence length. For the longest tested sequence (853,910nt) at *b* = *W* = 100, CapR required 514s and 1.65GB, whereas LinearCapR completed 610s and 10.6GB. Memory demands exceeded 120GB for *b* ≥ 200, making memory the practical limiting factor for long RNAs.

### 3.2 Accuracy

To assess the discriminative power of the structural-context probabilities, we used the bpRNA-1m(90) with pseudoknot-free reference structures. For each nucleotide i and context *δ* ∈ {*S, H, B, I, M, E* }, the probability *p* (*i, δ*) was treated as a one-vs-rest classifier score, and ROC curves and AUC values were computed (Fig.3).

**Figure 3.**
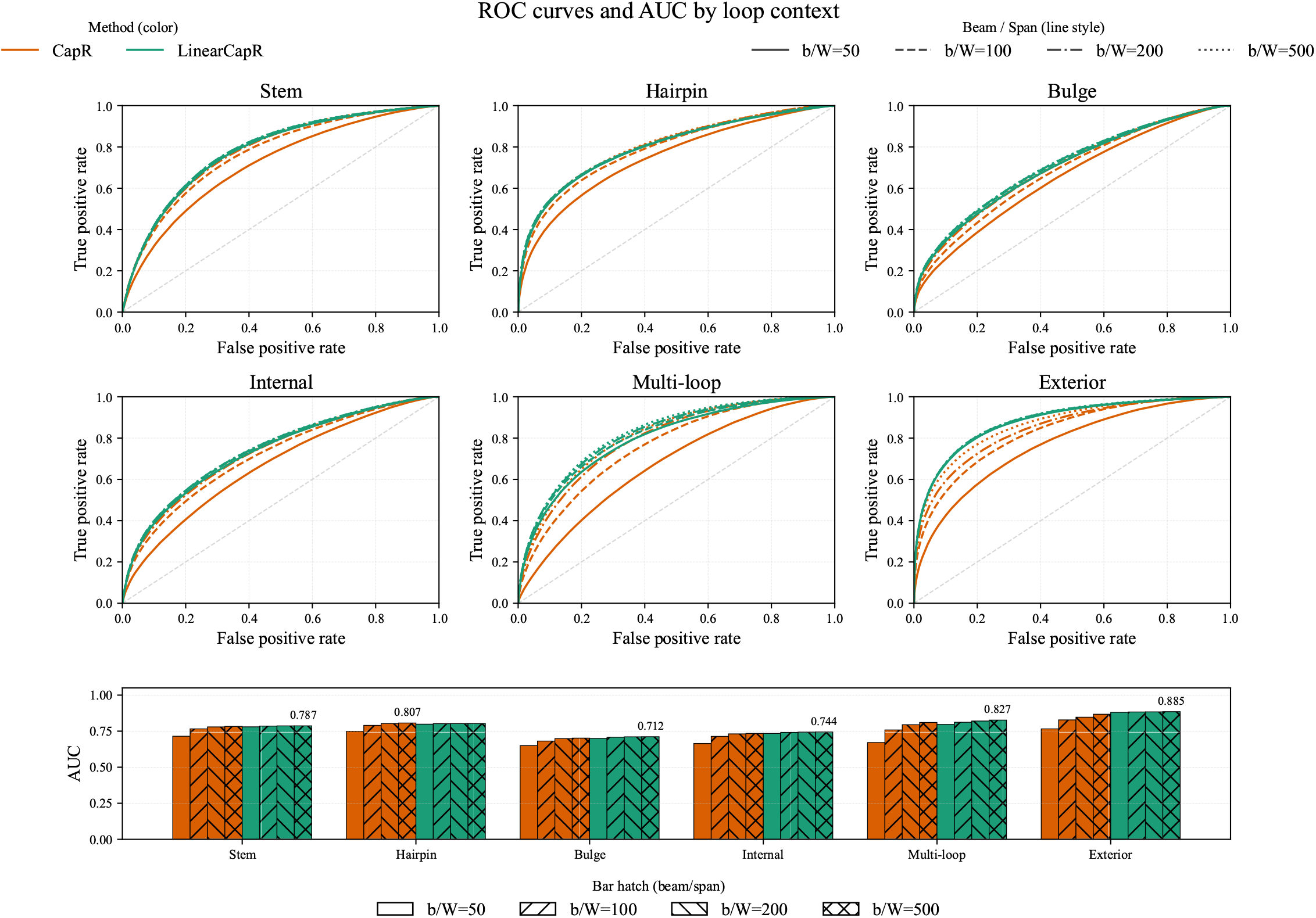
ROC curves for stems (S), hairpins (H), bulges (B), internal loops (I), multiloop (M), and exterior loops (E) comparing LinearCapR and CapR on bpRNA. Bars summarise AUC per context. Larger beam/window sizes improve ROC/AUC for both tools. Exterior loops—strongly influenced by long-range pairs—show the clearest LinearCapR advantage, consistent with its span-unrestricted search.

With *b* = *W* = 100, LinearCapR improved AUCs across all six contexts, from +0.012 (hairpins) to +0.055 (exterior), where long-range interactions are dominant. Improvements plateaued beyond *b* ≥ 200. Default parameter comparisons (CapR:

Turner1999; LinearCapR: Turner2004) are shown in the main text: Supplementary Fig. S2 provides matched Turner1999 results, where AUCs remain comparable overall, with LinearCapR favoured at small beams and CapR slightly higher at *b* = *W* = 300.

At *b* = *W* = 100, ROC curves for LinearCapR dominate those of CapR for every class, yielding AUC gains of 0.019 (stem), 0.012 (hairpin), 0.026 (bulge), 0.027 (internal), 0.053 (multiloop), and 0.055 (exterior). Increasing *b* from 100 to 200 yields *<* 0.003 macro-AUC improvement, indicating that *b* = 200 serves as a practical saturation point for bpRNA-1m(90). Whereas CapR requires spans *W >* 300 to reduce the gap for exterior and multiloop contexts, LinearCapR reaches comparable performance at *b* = 50.

To directly examine long-range stems, we computed AUC using only stems with pairing distance ≥ *L* ∈ { 150, 300} (Fig.4). LinearCapR showed substantial gains: at *b* = *W* = 100, AUC increased from 0.617 to 0.716 (*L* = 150) and from 0.633 to 0.714 (*L* = 300). Because CapR cannot represent stems longer than *W*, its performance decreases sharply under these criteria.

**Figure 4.**
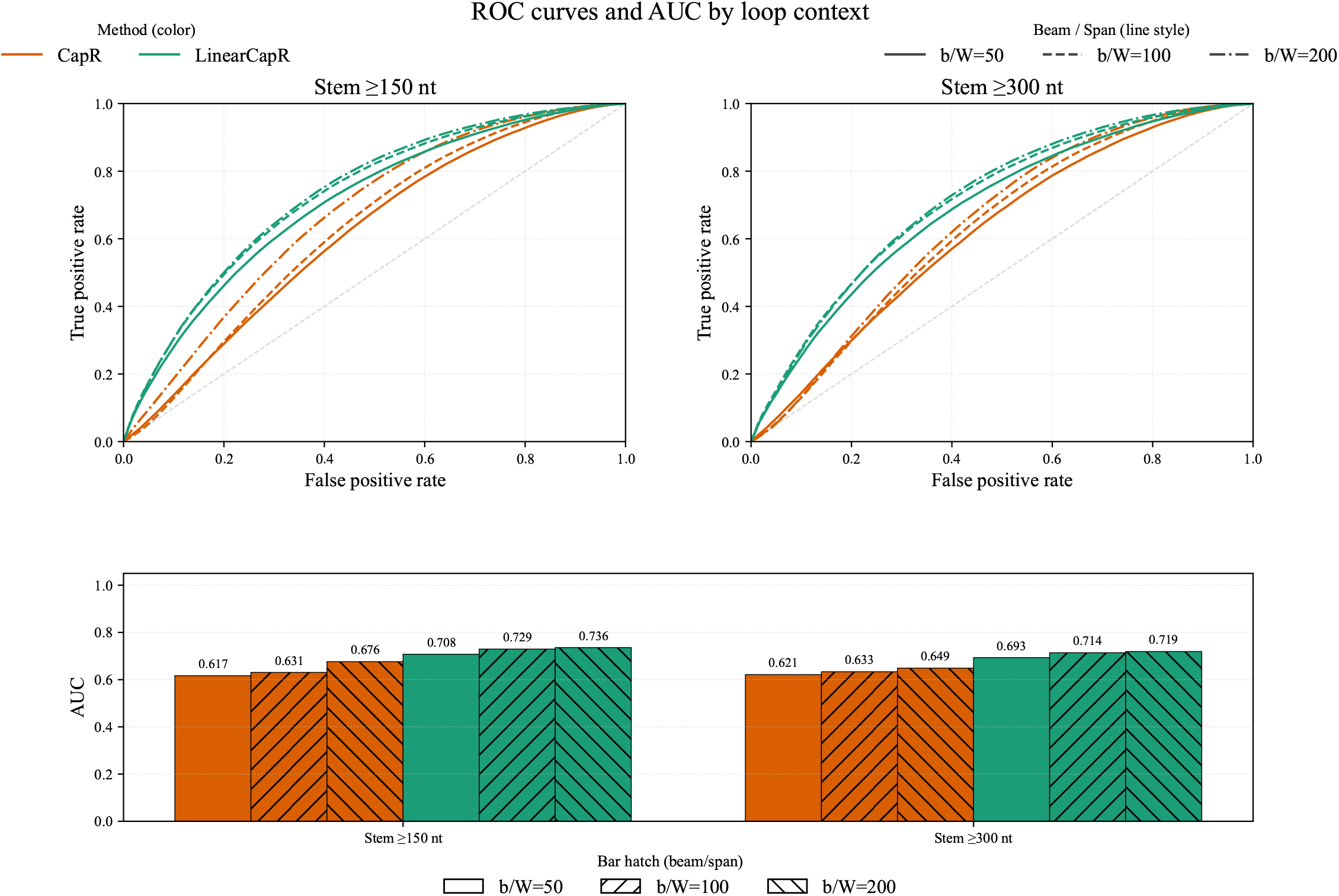
Long-range stem ROC curves and AUC for minimum distances of 150 and 300 nt. Even with a small beam (*b* = 50), LinearCapR surpasses CapR at *W* = 200 for these distant stems.

We also evaluated predicted context distributions on the RNAcentral panel. Although reference structures are unavailable, context entropy remained stable as *N* increased for LinearCapR, while decreasing under span-limited modeling (Supplementary Fig. S3), indicating that LinearCapR preserves ensemble diversity more faithfully for extremely long RNAs.

### 3.3 Beam-width dependence on ensemble quantities

Finally, we examined how beam width affects both the quality of ensemble estimates and computational cost. We performed beam-width sweeps on the 29,903 nt SARS-CoV-2 genome for *b* ranging from 5 to 200, under both Turner2004 and Turner1999 parameters (Fig. 5). The ensemble free energy *G*_ensemble_ in Turner2004 declines noticeably when *b <* 50 but changes little once *b* ≥ 100; Turner1999 shows the same flattening despite its different absolute values. Runtime and memory scaling for this sweep are provided in Supplementary Fig. S4, and dataset statistics in Supplementary Table S1.

**Figure 5.**
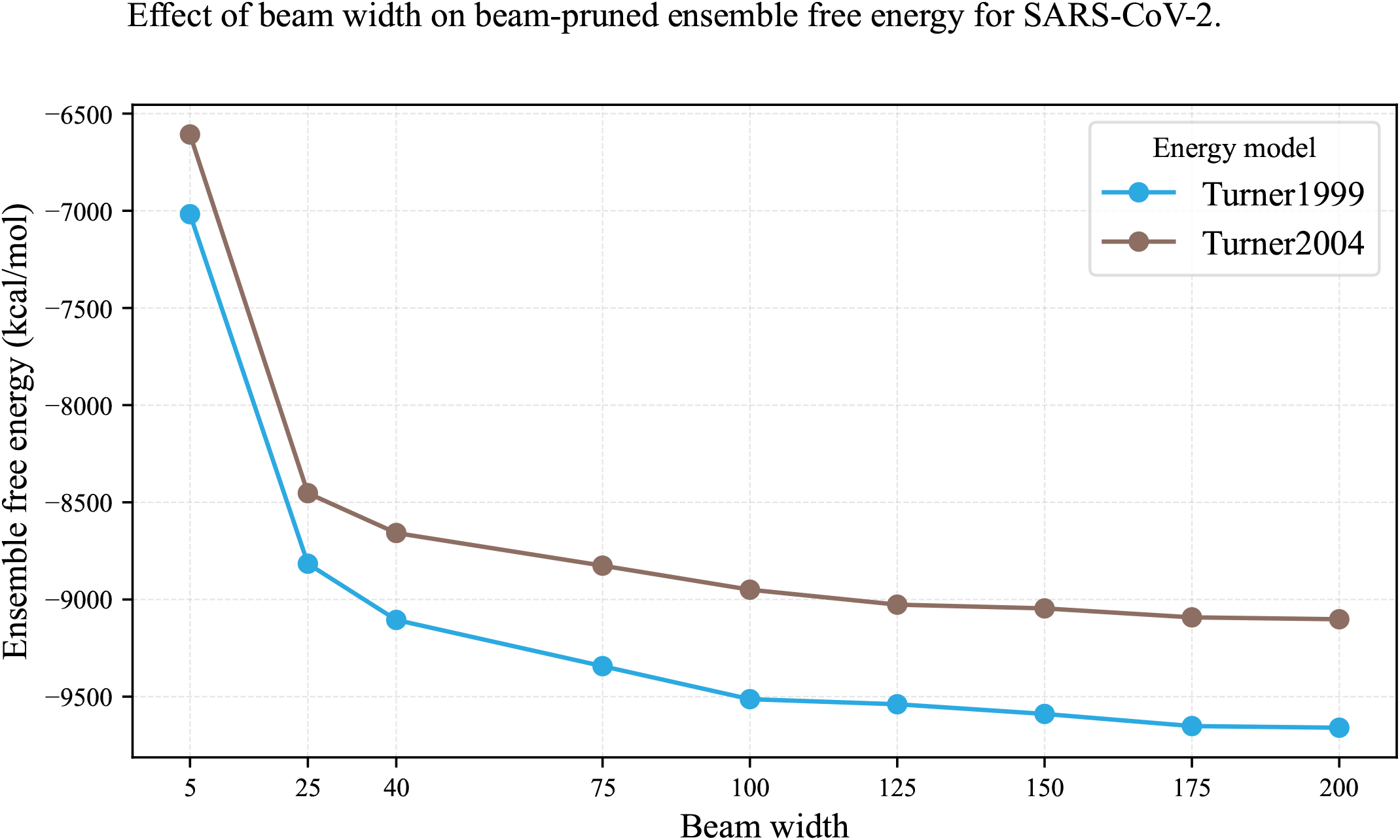
Effect of beam width on ensemble free energy *G*_ensemble_ for the SARS-CoV-2 genome. Turner2004 (brown) and Turner1999 (sky blue) curves are overlaid; the latter yield different absolute energies because of the parameters, but both models flatten beyond *b* ≈ 100. Runtime and memory trajectories are shown in Supplementary Fig. S4.

These results indicate that relatively moderate beam width *b* ≥ 100 already stabilizes ensemble estimates for genome-scale RNAs.

## 4 Discussion

LinearCapR is the first method to compute CapR-style RNA structural-context posterior probabilities in linear time while retaining long-range base-pair interactions. Eliminating the base-pair span limit substantially improves the characterization of exterior and multiloop contexts and enhances detection of long-range stems, which are prevalent in viral RNAs and long noncoding RNAs. Table 2 summarizes key implementation differences between CapR and LinearCapR, including search limits, the time and space complexity, and supported energy parameters.

**Table 2:** Key implementation differences to guide parameter choice.

| Aspect | CapR | LinearCapR |
| --- | --- | --- |
| Search limit | Span limit $W$ | Beam width $b$ (no span cap) |
| Complexity | $O(NW^2)$ | $O(Nb^2 + NbC^2)$ |
| Energies | Turner1999 | Turner2004 / Turner1999 |

The primary cost of beam pruning is increased memory usage. Sections 3.1–3.3 collectively suggest that beam widths in the range of 100–200 offer a practical trade-off among accuracy, runtime, and memory for transcriptome- and genome-scale RNAs. On the SARS-CoV-2 genome, ensemble thermodynamic estimates stabilize once *b* ≥ 100, and RNAcentral experiments demonstrate that LinearCapR avoids the bias toward local structures that arises when long-range stems are forcibly truncated.

Future directions include compressed state encodings, GPU parallelization, and adaptive beam control to reduce peak memory demands. Integration with downstream analyses such as RBP-binding prediction or RNA-structure-aware functional annotations represents another promising path.

Overall, LinearCapR enables scalable and biologically meaningful structural profiling of long RNAs with ensemble fidelity. CapR-style context profiles have been shown to improve models of RNA-binding protein specificity from CLIP-seq data. Because LinearCapR scales these profiles to transcriptome- and viral genome-length RNAs without span constraints, it enables systematic re-analysis of existing CLIP datasets and joint modelling of sequence and context preferences. We leave such RBP-focused applications to future work.

### Limitations

LinearCapR inherits several modelling assumptions from prior SCFG-based RNA folding methods. First, we restrict our search space to pseudoknot-free secondary structures, and non-canonical base pairs are treated as unpaired residues. Second, the grammar bounds contiguous unpaired runs to *C* = 30 nucleotides; this covers 99.57% of multiloop segments in bpRNA (Supplementary Fig. S1) but may underrepresent exceptionally long loops. Third, the beam-pruned inside-outside algorithm yields an approximation of the true Boltzmann ensemble. Our experiments suggest that moderate beam widths (*b* ≈ 100-200) are sufficient for stable ensemble estimates.

## Supplementary data

Supplementary data include all the graphs and tables, and are available at github: https://github.com/TakumiOtagaki/LinearCapR_ComputationalExperiments. The dataset summary is also available, so every figure in the main text can be regenerated from these artifacts hosted in the public repository.

## 5 Acknowledgements

This research was conducted using the FUJITSU Supercomputer PRIMEHPC FX1000 and FUJITSU Server PRIMERGY GX2570 (Wisteria/BDEC-01) at the Information Technology Center, The University of Tokyo.

