## Supplementary Information for "LinearCapR: Linear-time computation of per-nucleotide structural-context probabilities of RNA without base-pair span limits"

### Algorithm S1: Detailed algorithms

Here $Z$ denotes the partition function estimated by the Inside pass. The auxiliary terms $B_{\mathrm{Hairpin}}$, $B_{\mathrm{Loop}}$, and $B_{\mathrm{MultiUnpaired}}$ accumulate the Boltzmann-weighted contributions from hairpins, bulge/internal loops, and multibranch-loop unpaired segments, respectively; prefix-sum buffers are used in the multibranch case to aggregate contiguous updates efficiently. The repeated range additions to the structural profile (e.g., $p([i,j],H)\ \mathrel{+}= \ldots$) are implemented via 1D difference-array buffers that record the contributions in $O(1)$ time per range update at the range boundaries and materialize the prefix sum only once per context. This keeps the profile aggregation linear-time despite the nested loops over $p$, $q$, and running unpaired segments. This algorithm shows the full Outside recursion that complements Algorithm 1 in the main text.

**Algorithm S1.1** Outside algorithm in LinearCapR

**procedure** CALC_OUTSIDE
  **for** $j = N \ldots 1$ **do**
    ▷ Outer
    $\beta_{\mathrm{Outer}}(j) \mathrel{+}= \beta_{\mathrm{Outer}}(j+1) \cdot t(\mathrm{Outer} \to \mathrm{Outer})$
    ▷ Stem
    **for all** $[i,j] \in \alpha_{\mathrm{Stem}}$ **do**
      $\beta_{\mathrm{Outer}}(i) \mathrel{+}= \beta_{\mathrm{Outer}}(j+1) \cdot \alpha_{\mathrm{Stem}}(i,j) \cdot t(\mathrm{Outer} \to \mathrm{Outer} \cdot \mathrm{Stem})$
    ▷ StemEnd
    **for all** $[i,j] \in \alpha_{\mathrm{StemEnd}}$ **do**
      $\beta_{\mathrm{StemEnd}}(i,j) \mathrel{+}= \beta_{\mathrm{Stem}}(i-1,j+1) \cdot t(\mathrm{Stem} \to \mathrm{StemEnd})$
    ▷ Multi
    **for all** $[i,j] \in \alpha_{\mathrm{Multi}}$ **do**
      $\beta_{\mathrm{Multi}}(i,j) \mathrel{+}= \beta_{\mathrm{StemEnd}}(i,j) \cdot t(\mathrm{StemEnd} \to \mathrm{Multi})$
    ▷ MultiBif
    **for all** $[i,j] \in \alpha_{\mathrm{MultiBif}}$ **do**
      $\beta_{\mathrm{MultiBif}}(i,j) \mathrel{+}= \beta_{\mathrm{Multi1}}(i,j) \cdot t(\mathrm{Multi1} \to \mathrm{MultiBif})$
      **for** $n = 0 \ldots C$ **do**
        $\beta_{\mathrm{MultiBif}}(i,j) \mathrel{+}= \beta_{\mathrm{Multi}}(i-n,j) \cdot t(\mathrm{Multi} \to \mathrm{MultiBif})$
    ▷ Multi2
    **for all** $[i,j] \in \alpha_{\mathrm{Multi2}}$ **do**
      $\beta_{\mathrm{Multi2}}(i,j) \mathrel{+}= \beta_{\mathrm{Multi1}}(i,j) \cdot t(\mathrm{Multi1} \to \mathrm{Multi2})$
    **for all** $[k,i-1] \in \alpha_{\mathrm{Multi1}}$ **do**
      $\beta_{\mathrm{Multi1}}(k,i-1) \mathrel{+}= \alpha_{\mathrm{Multi2}}(i,j) \cdot \beta_{\mathrm{MultiBif}}(k,j) \cdot t(\mathrm{MultiBif} \to \mathrm{Multi1} \cdot \mathrm{Multi2})$
      $\beta_{\mathrm{Multi2}}(i,j) \mathrel{+}= \alpha_{\mathrm{Multi1}}(k,i-1) \cdot \beta_{\mathrm{MultiBif}}(k,j) \cdot t(\mathrm{MultiBif} \to \mathrm{Multi1} \cdot \mathrm{Multi2})$
    ▷ Stem
    **for all** $[i,j] \in \alpha_{\mathrm{Stem}}$ **do**
      $\beta_{\mathrm{Stem}}(i,j) \mathrel{+}= \alpha_{\mathrm{Outer}}(i-1) \cdot \beta_{\mathrm{Outer}}(j+1) \cdot t(\mathrm{Outer} \to \mathrm{Outer} \cdot \mathrm{Stem})$
      $\beta_{\mathrm{Stem}}(i,j) \mathrel{+}= \beta_{\mathrm{Stem}}(i-1,j+1) \cdot t(\mathrm{Stem} \to \mathrm{Stem})$
      **for** $n = 0 \ldots C$ **do**
        $\beta_{\mathrm{Stem}}(i,j) \mathrel{+}= \beta_{\mathrm{Multi2}}(i,j+n) \cdot t(\mathrm{Multi2} \to \mathrm{Stem})$
      **for all** $p \le i < j \le q,\ 0 < (i-p)+(q-j) \le C$ **do**
        $\beta_{\mathrm{Stem}}(i,j) \mathrel{+}= \beta_{\mathrm{StemEnd}}(p,q) \cdot t(\mathrm{StemEnd} \to \mathrm{Stem})$

**Algorithm S1.2** Full calculation of structure profile

**procedure** CALC_PROFILE
- **for all** $[i,j] \in \alpha_{\text{StemEnd}}$ **do**
  - $p([i,j],H) \mathrel{+}= \frac{1}{Z}\,\beta_{\text{StemEnd}}(i,j) \cdot B_{\text{Hairpin}}(i-1,j+1)$
  - **for** $p = i+1 \ldots \min(i+C,\, j-1)$ **do**
    - $p([\,i,p-1\,],B) \mathrel{+}= \frac{1}{Z}\,\alpha_{\text{Stem}}(p,j) \cdot \beta_{\text{StemEnd}}(i,j) \cdot B_{\text{Loop}}(i-1,\, j+1,\, p,\, j)$
  - **for** $q = \max(j-C,\, i+1) \ldots j-1$ **do**
    - $p([\,q+1,j\,],B) \mathrel{+}= \frac{1}{Z}\,\alpha_{\text{Stem}}(i,q) \cdot \beta_{\text{StemEnd}}(i,j) \cdot B_{\text{Loop}}(i-1,\, j+1,\, i,\, q)$
  - **for** $p = i+1 \ldots \min(i+C,\, j-1)$ **do**
    - **for** $q = \max(p-i+j-C,\, p+1) \ldots j-1$ **do**
      - $p([\,i,p-1\,],I) \mathrel{+}= \frac{1}{Z}\,\alpha_{\text{Stem}}(p,q) \cdot \beta_{\text{StemEnd}}(i,j) \cdot B_{\text{Loop}}(i-1,\, j+1,\, p,\, q)$
      - $p([\,q+1,j\,],I) \mathrel{+}= \frac{1}{Z}\,\alpha_{\text{Stem}}(p,q) \cdot \beta_{\text{StemEnd}}(i,j) \cdot B_{\text{Loop}}(i-1,\, j+1,\, p,\, q)$
- **for all** $[p,j] \in \alpha_{\text{MultiBif}}$ **do**
  - **for** $i = \max(1,\, p-C) \ldots p-1$ **do**
    - $p([\,i,p-1\,],M) \mathrel{+}= \frac{1}{Z}\,\alpha_{\text{MultiBif}}(p,j) \cdot \beta_{\text{Multi}}(i,j) \cdot B_{\text{MultiUnpaired}}(i,\, p-1)$
- **for all** $[i,q] \in \alpha_{\text{Stem}}$ **do**
  - **for** $j = q+1 \ldots \min(N,\, q+C)$ **do**
    - $p([\,q+1,j\,],M) \mathrel{+}= \frac{1}{Z}\,\alpha_{\text{Stem}}(i,q) \cdot \beta_{\text{Multi2}}(i,j) \cdot B_{\text{MultiUnpaired}}(q+1,\, j)$
- **for all** $[i,j] \in \alpha_{\text{Stem}}$ **do**
  - $p(i,S) \mathrel{+}= \frac{1}{Z}\,\alpha_{\text{Stem}}(i,j) \cdot \beta_{\text{Stem}}(i,j)$
  - $p(j,S) \mathrel{+}= \frac{1}{Z}\,\alpha_{\text{Stem}}(i,j) \cdot \beta_{\text{Stem}}(i,j)$
- $p(i,E) = \frac{1}{Z}\,\alpha_{\text{Outer}}(i-1) \cdot \beta_{\text{Outer}}(i+1)$

**Fig. S1: Multiloop unpaired run lengths**

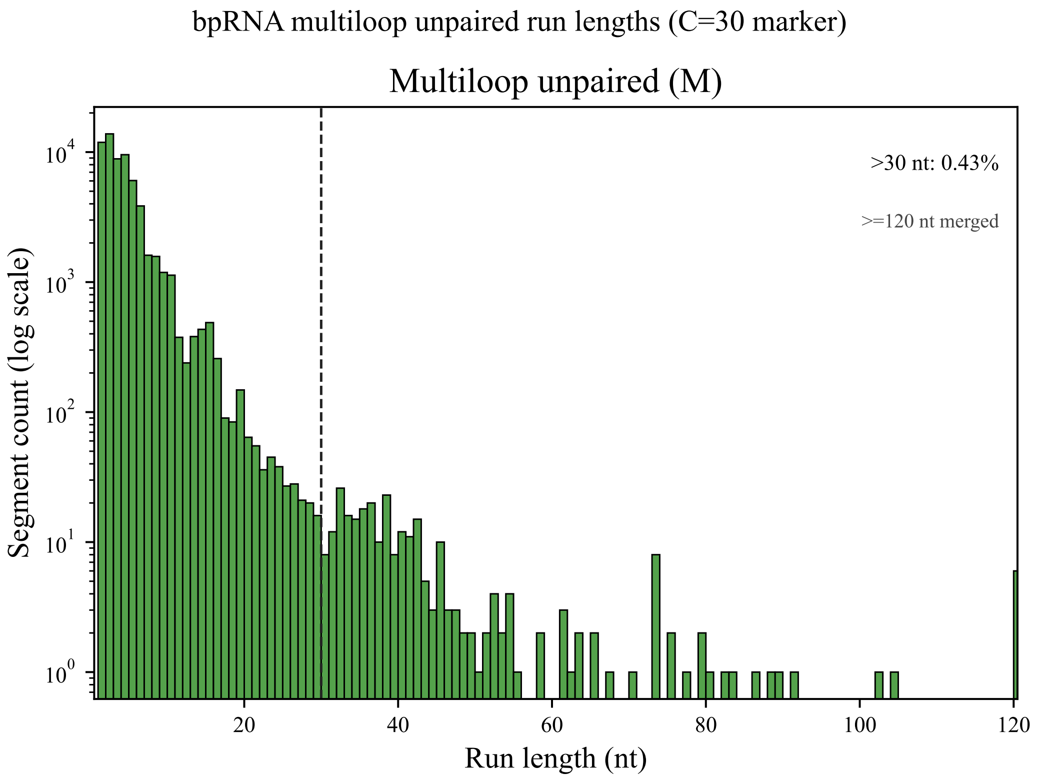


Fig. S1: Distribution of consecutive unpaired nucleotides within multiloops in bpRNA-1m(90). The $y$-axis is logarithmic. Only 0.43% of runs exceed 30 nt, indicating that the $C = 30$ cap covers 99.57% of multiloop unpaired segments.

### Table S1: Benchmark datasets

**Table S1.** Overview of benchmark datasets considered in this study. Counts and lengths follow the summaries produced by the plotting pipeline (`https://github.com/TakumiOtagaki/LinearCapR_ComputationalExperiments/graphs/dataset_summary/dataset_summary.csv`).

| Dataset | Sequences | Length range | Role in study |
|---|---|---|---|
| bpRNA-1m(90) subset | 24,901 | 11–4,065 nt | Accuracy and runtime comparison |
| RNAcentral long RNAs | 20 | 4,793–853,910 nt | Scalability assessment |
| SARS-CoV-2 genome | 1 | 29,903 nt | Beam width sensitivity |

### Fig. S2: Accuracy with Turner1999 parameters

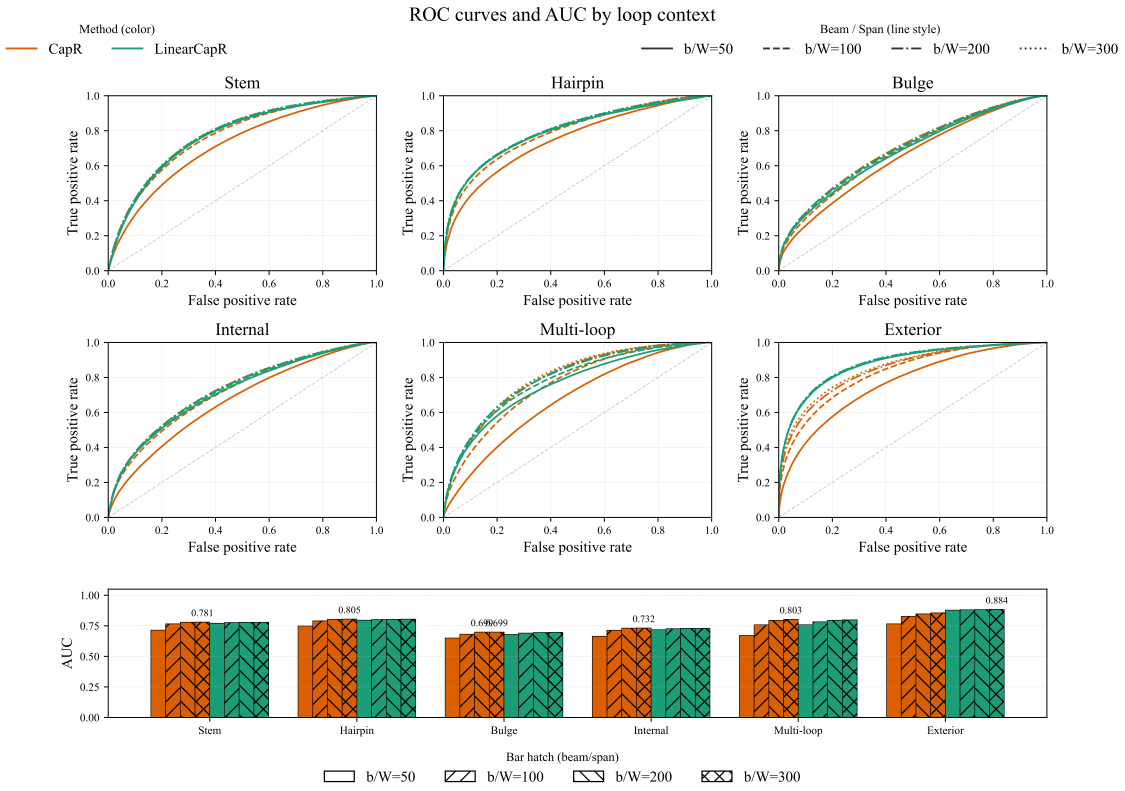


Fig. S2: ROC curves and AUC when both CapR and LinearCapR use Turner1999. LinearCapR matches or exceeds CapR for several contexts, especially at smaller beam widths; when the beam/window increases to 300, CapR is slightly higher for many contexts. For exterior loops, LinearCapR yields higher AUC than CapR across all tested beam/window sizes.

### Fig. S3: Long-range stems with Turner1999 parameters

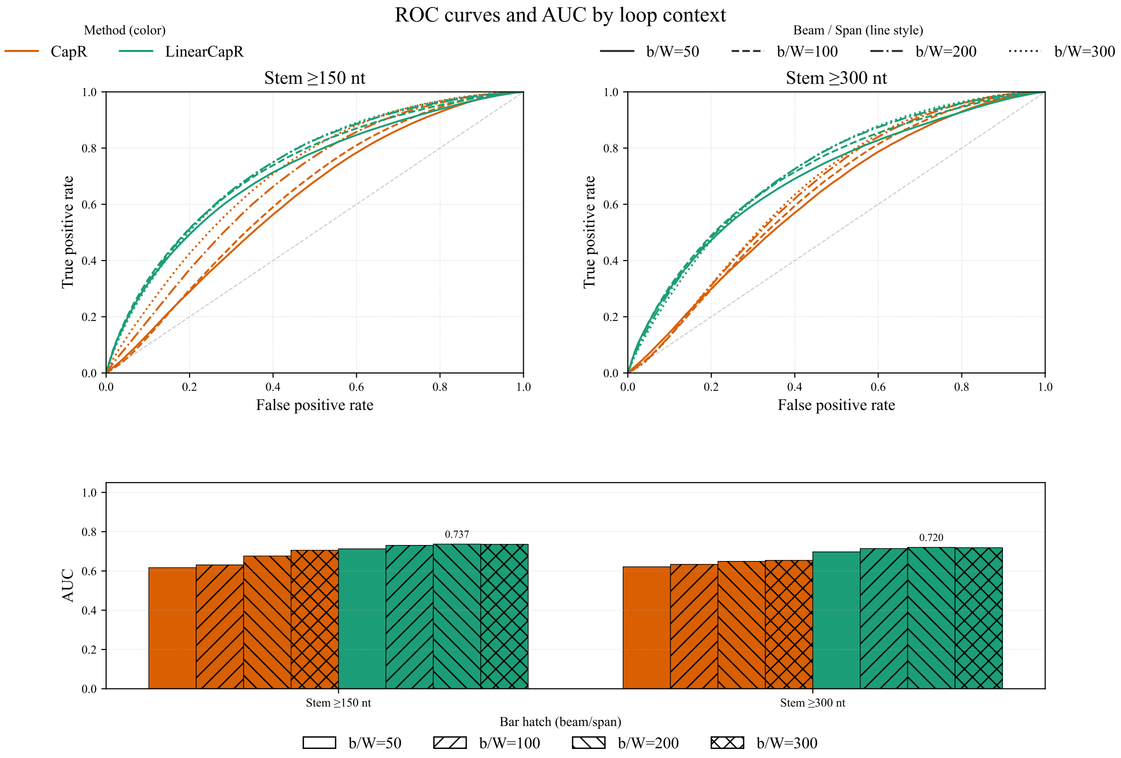


Fig. S3: ROC curves and AUC for long-range stems (minimum distances 150 nt and 300 nt) when both CapR and LinearCapR use Turner1999. LinearCapR generally attains higher AUCs across beam/window settings for these distant stems.

### Fig. S4: Beam-sweep runtime and memory

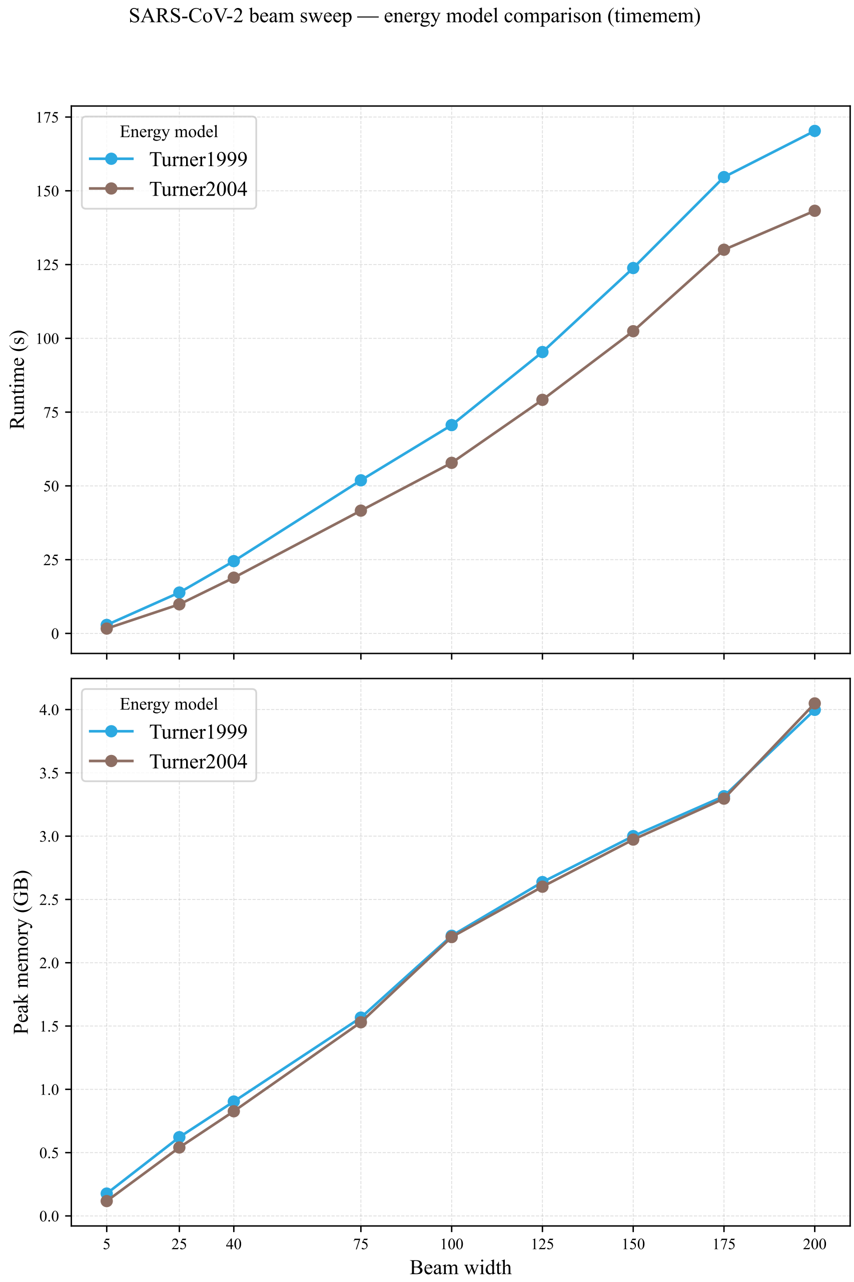


Fig. S4: Runtime and peak memory versus beam width for the SARS-CoV-2 genome under Turner2004 (sky blue) and Turner1999 (brown). Both models scale roughly linearly with $b$; Turner1999 runs longer but uses similar memory.

### Text S2: RNAcentral sampling protocol

To assemble the 20 long-RNA test cases in RNAcentral we used `LinCapR_Experiments/scripts/data_preprocessing/sample_log_bins_from_tsv.py`. The script reads a tab-separated file with header `name<TAB>length`, builds 20 log-spaced bins between the minimum and maximum lengths, and samples one entry per bin with a fixed random seed (`--seed 42`, `--k 1`). Bin edges are computed on the log scale and each bin covers $[\,\mathrm{edge}_i, \mathrm{edge}_{i+1} - 1\,]$. The resulting list of accession IDs is written to `--out` and used to fetch the sequences; summary statistics are exported as `graphs/dataset_summary/dataset_summary.csv` to reproduce the counts and length ranges reported in the main text.
